# SpikeCleaner: An Algorithm to Label Unit Quality After Automated Spike Sorting

**DOI:** 10.64898/2026.06.18.733033

**Authors:** Diksha Zutshi, Daniil Berezhnoi, Anjesh Ghimire, Jeremiah Hartner, David S. Kim, Brendon O. Watson

## Abstract

**Gap:** Automated spike sorting algorithms have revolutionized the way neuronal activity is extracted from extracellular recordings, yet they remain imperfect. Specifically, inaccurate acceptance of noise-based units not only leaves researchers with clusters that require extensive manual curation, an essential but time-consuming process, that also leads to significant subjectivity in the selection of units. In an era of high-density probes like Neuropixels, where an hour of data can exceed 80 GB, manual curation is no longer scalable, automation of standard criteria can speed data curation and ensure quality of datasets. Here, we developed a semi-automated curation pipeline to label the quality of units after automated curation by Kilosort.

**Approach:** Our algorithm standardizes criteria for labeling of Noise, Multi-Unit Activity (MUA), and Good Units using a combination of spike rate, spike timing metrics (from autocorrelogram), and waveform-based physiological features such as peak amplitude, slopes, half-width, and inter-channel correlation. Based on these features, clusters are assigned standardized labels (good, noise, multi-unit activity) that can be imported directly into Phy, where they serve as curation aids rather than absolute classifications, supporting but not replacing expert judgment. Heuristically, “noise” units are those unlikely to be neuronal in origin; “MUA” includes units with significant neural contribution (i.e., neuronal waveform) but with some degree of clear imperfection to be further cleaned, and “good” units are those without any clear deviation from ideal unit criteria. By ensuring accurate selection of acceptable units, we enable robust downstream analyses such as neural decoding and longitudinal tracking of neuron identity. Thresholds for all metrics were chosen to maximize the matching of algorithm output to that of 2 expert manual curators. Of note, users may alter thresholds either based on their own judgment or using an included tool to semi-automatically find thresholds that optimize SpikeCleaner with their own expert curation.

**Results:** To benchmark, we compared the outputs of our algorithm to expert-labels curated in Phy by two expert users across three recordings. SpikeCleaner achieved an average of **97% accuracy vs. experts & 92% F1 score** in classifying **Single Units**. It achieved an accuracy of **97% & 92% F1 score** in full-category agreement (SU, MUA, Noise), and **97% accuracy & 95% F1 score** in distinguishing **Neuronal vs. Non-Neuronal** units.

## INTRODUCTION

Spike sorting is crucial for recording the activity of multiple individually-tracked neurons in a local population to study network dynamics. A number of state-of-the-art spike sorting algorithms analyze voltage recordings from physically adjacent electrodes to provide an initial sorting of spikes into units or putative neurons.^**1-5**^ However, despite significant advances in these automated spike sorting algorithms, current tools remain prone to errors, particularly when handling noisy data or complex spiking activity.^**15**^ These errors require downstream manual “curation” of the resultant spike sorting assignments by human users, requiring tens of minutes to hours of work for single recordings. Specifically, this human-executed manual review and refinement of the output of automated spike sorting algorithms attempts to ensure that each detected cluster of spikes truly corresponds to a single neuron. Manual curation tools, such as Phy,^**6**^ are widely adopted for curation tasks, including labeling the quality of units as noise, good or multi-unit. However, Phy and similar tools function primarily as graphical user interfaces without integrated algorithmic decision-making. They therefore rely completely on the user’s judgment, leading to the consumption of many hours of human time and to subjective, non-standardized results. Moreover, due to the scale of high-density datasets (e.g., 1 hour Neuropixels 1.0 or 2.0 probe recording, with their 384 channels sampled at 30 kHz and with 2 bytes per sample, will produce over ∼80 GB of unprocessed raw data), exhaustive manual curation becomes increasingly infeasible.^**7**^ Additionally, rigorous manual curation of longer data sets leads to fatigue, further adding bias and reducing reproducibility. Even without fatigue, different users may apply different criteria—leading to heterogeneity. Finally, the lack of recorded and reportable objective criteria leads to near impossibility of reproduction of findings even from the same recordings.

As Buccino et al.^**6**^, decisions around cluster merging or deletion are influenced by the curator’s experience, the dataset type, and even the specific brain region or neuronal subtype involved. This inconsistency introduces irreproducibility in downstream neural analyses and complicates benchmarking across studies.^**7**^ Thus, basic quality control could achieve homogenization across users, save time, and enable faster turnaround time for initial iterative parameter testing.

To address these challenges, we developed SpikeCleaner, an open-source MATLAB-based algorithm for threshold-driven, automated neural cluster curation, achieving performance comparable to expert-level accuracy. Because SpikeCleaner relies on explicitly defined, hard-coded thresholds rather than data-specific training or model fitting, its performance can be automatically reproduced and applied across diverse datasets, electrode configurations, and experimental conditions. Furthermore, thresholds used in SpikeCleaner can be recorded and reported as methods. Rather than replacing expert judgment, SpikeCleaner is designed to complement it by enabling side-by-side comparison between algorithmic and manual curation, both visually in Phy and quantitatively through summary metrics. This dual evaluation framework enables users to fine-tune curation parameters based on their data characteristics and individual curation style. Finally, while we provide default thresholds, we also provide tools for users to determine new thresholds in a semi-automated manner for their own situation or user group.

## METHODS

### Overall Workflow

SpikeCleaner operates immediately downstream of Kilosort and upstream of Phy (**Figure 1**). It uses spike times, cluster assignments, and channel information from Kilosort as inputs while adding an additional layer of biologically informed curation and outputting results to Phy-readable files. Whereas Kilosort performs initial spike detection and clustering based on template matching, SpikeCleaner reevaluates spikes from each resulting cluster using waveform shape, firing statistics, spatial correlations, and autocorrelogram structure to assign standardized labels (Noise, MUA, or Single Unit). As illustrated in Figure 1, SpikeCleaner acts as an intermediate quality-control module between Kilosort and Phy, automating and standardizing the most error-prone and subjective components of manual curation while preserving full compatibility with existing workflows.

**Figure 1.**
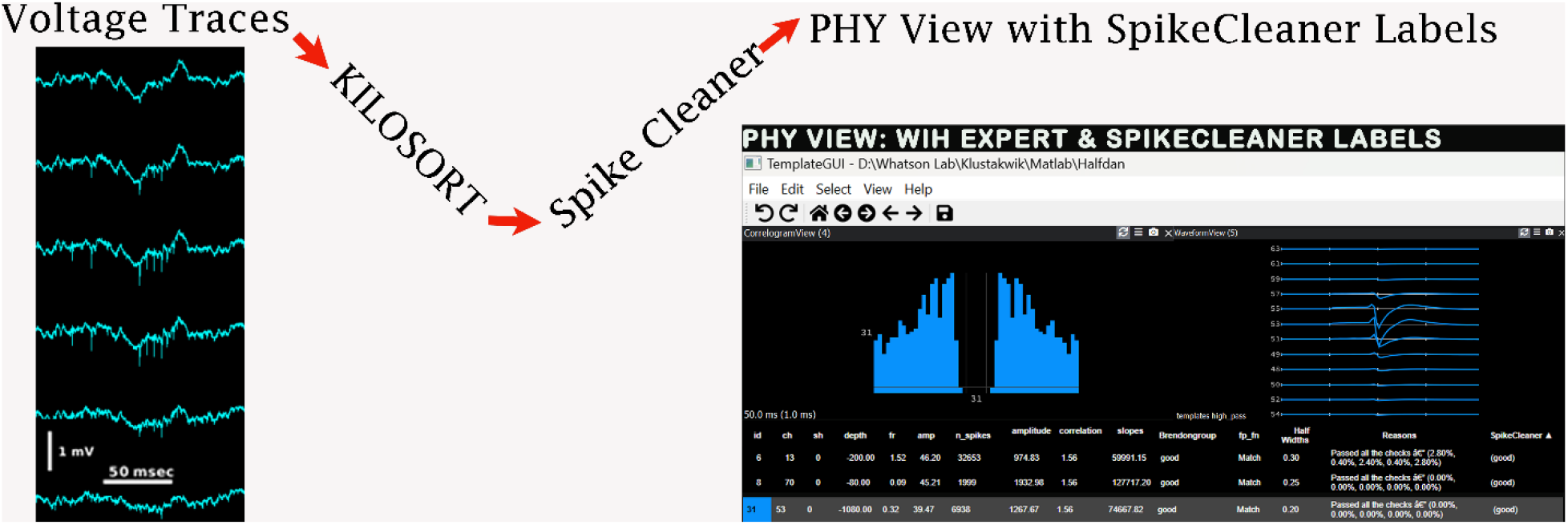
Sequence of events

SpikeCleaner performs standardized extraction and processing of spike waveforms and spike times to ensure reproducible downstream feature computation and biological quality assessment. The full pipeline is implemented in the core functions dz_Curate.m, dz_getWaveform.m, and dz_filterWaveform.m within the SpikeCleaner framework. For each recording, SpikeCleaner loads the outputs of Kilosort: spike_times.npy, spike_clusters.npy, channel_positions.npy, and rez.mat. These files contain spike timestamps (in samples), the associated cluster ID for each spike, the sampling rate, and the active channels used during spike sorting. After loading, spikes are grouped by cluster, and a representative set of waveform snippets is extracted for each cluster.

### Data Preprocessing

SpikeCleaner extracts raw spike waveforms using dz_getWaveform.m. For each spike, a window of ±2 ms around the spike time is taken. Sampling rate is extracted from params.py, which is an output of Kilosort ^**1**^. For a 20 kHz sampling rate, this corresponds to ∼80 samples per spike. Waveforms are extracted across all active channels used for spike sorting. This multi-channel representation preserves spatial information needed to identify the best channel, detect biological waveform shapes, and compute inter-channel correlations used for noise rejection. If a cluster exceeds 10,000 spikes, a random 10% subset is used to reduce memory and computational load while maintaining representative waveform statistics. For each cluster, we compute the across-spike means to obtain a single average multichannel waveform ([time × channels]). This reduces noise and provides a stable template for downstream feature extraction.

Extracted waveforms are then filtered using dz_filterWaveform.m. First, each channel undergoes a zero-phase infinite impulse response Butterworth high-pass filter at 150 Hz (3rd order, filtfilt.m in MATLAB), matching Phy’s preprocessing and removing low-frequency drift while preserving spike morphology. After high-pass filtering, a moving-average smoothening window of 9 samples is applied per channel to stabilize the global waveform shape and reduce residual high-frequency fluctuations. Importantly, the smoothed waveforms are used only for identifying the overall spike shape, whereas all quantitative decisions: amplitude, slope, half-width, and inter-channel correlation are computed exclusively on the high-passed (unsmoothed) waveforms to avoid altering biologically relevant features.

### Data Structures & Extraction

SpikeCleaner processes data cluster-by-cluster inside a loop. To keep the pipeline fast and reproducible, we extract expensive-to-compute objects once, store them in standardized MATLAB data structures, and save them as .mat files that can be reloaded in later loop iterations (or future reruns) without re-reading the raw .dat.

Before waveform extraction begins, the pipeline prompts the user to specify the data format:

1. If the data is already in units of voltage, waveform extraction proceeds directly.
2. If the data is in binary (.dat) format, as is typical for Intan RHD recordings, the data is stored as int16 ADC counts.

In case of binary data recorded via Intan:

*extractedData = 0.195 * (extractedData - 32768); % units = microvolts* ; *conversion from int16 to uV*

This ensures that all waveform-derived features (amplitude, slope, half-width, correlation) are computed in physically meaningful voltage units rather than arbitrary ADC counts.

This ±2 ms around the spike time is saved as the waveform snippet in the **Output** folder as animalname.mat file within the data folder by **dz_getWaveform()**. Inside this file, the waveforms are organized in a cell array named wf, where each cell corresponds to a cluster ID. Thus, wf{cluster_id} contains the waveform snippets for that cluster, stored as a matrix of [number of samples × number of channels]. This structure allows flexible storage across clusters and enables efficient reuse of the extracted voltage-scaled waveforms in subsequent analysis loops without re-reading the raw .dat file.

This file is read by **dz_filterWaveform()**, which is called within **dz_Curate()**, which generates the animalname_filtered.mat file in the same folder as the unfiltered data. This file contains the filtered versions of the previously extracted ±2 ms waveform snippets and serves as the processed signal representation used for downstream feature extraction and quality control.

### The Standard/Default pipeline of SpikeCleaner if user provides none

The Journey to Acceptance goes from left to right and is a comprehensive set of quality checkpoints to assess cluster quality. Each of these tests are performed on features extracted from the waveforms or the ACG of the cluster. All clusters start with the assumption that they are Good/Single Unit and are discarded at different check points if they don’t meet the standard criterion of feature threshold. If a cluster does not pass the checks in the red box (steps A - G) it will be labelled as Noise, if it does not pass the check in yellow box (step H), it will be labelled as multi-unit activity (MUA). If a cluster passes all checks, it will be labelled as Good/Single Unit (green box/ step I). While the figure above is the default and recommended order of steps for SpikeCleaner, users can re-arrange the order of these steps as they best see fit.

#### 1. Firing Rate Analysis

As illustrated in **Figure 2-A**, SpikeCleaner, performs a low-firing-rate screen to eliminate clusters unlikely to represent biologically meaningful units. For each cluster, the algorithm computes its average firing rate using the true total number of spikes, not the 10% subsampled set used later for waveform extraction. This ensures that firing-rate estimates remain accurate and unaffected by downsampling. The firing rate is obtained by calling a subfunction in **dz_curate(), evaluateLowFiringRates(recordingDuration**,**firingThreshold)** from within the pipeline according to the order chosen by the user, which divides the full spike count by the recording duration, and clusters firing below the default threshold of 0.05 Hz (<1 spike every 20 seconds) are immediately flagged as Noise and excluded from subsequent waveform or ACG evaluation. This threshold reflects the fact that extremely sparse clusters are typically dominated by noise events, threshold crossings, or unstable detections rather than stable neuronal activity. SpikeCleaner also logs a human-readable reason for each rejected cluster (e.g., “Low firing rate: 0.012 Hz < 0.050 Hz”) to preserve transparency, while clusters that pass this screen proceed to the next stage of biological waveform assessment.^**9**^

~~~
*
For every loop run:
If ∼good(ix)
  continue;
end
i = uclu(ix); %uclu being the unique clusters
tst = ts(clu == i); %clu being the variable where we read spike_clusters.npy
totalSpikes=length(tst); %being the spike times of all the spikes in that particular cluster
Averagefiringrate=totalSpikes/recordingDuration;
if(Averagefiringrate<firingThreshold)
  noiseReasons{ix} = ‘lowRate’;
  Reasons{ix} = sprintf(‘Low firing rate: %.3f Hz < %.3f Hz (Threshold)’, Averagefiringrate, firingThreshold);
  good(ix)=false;
  continue;
else
  good(ix)=true;
end
*
~~~

**Figure 2.**
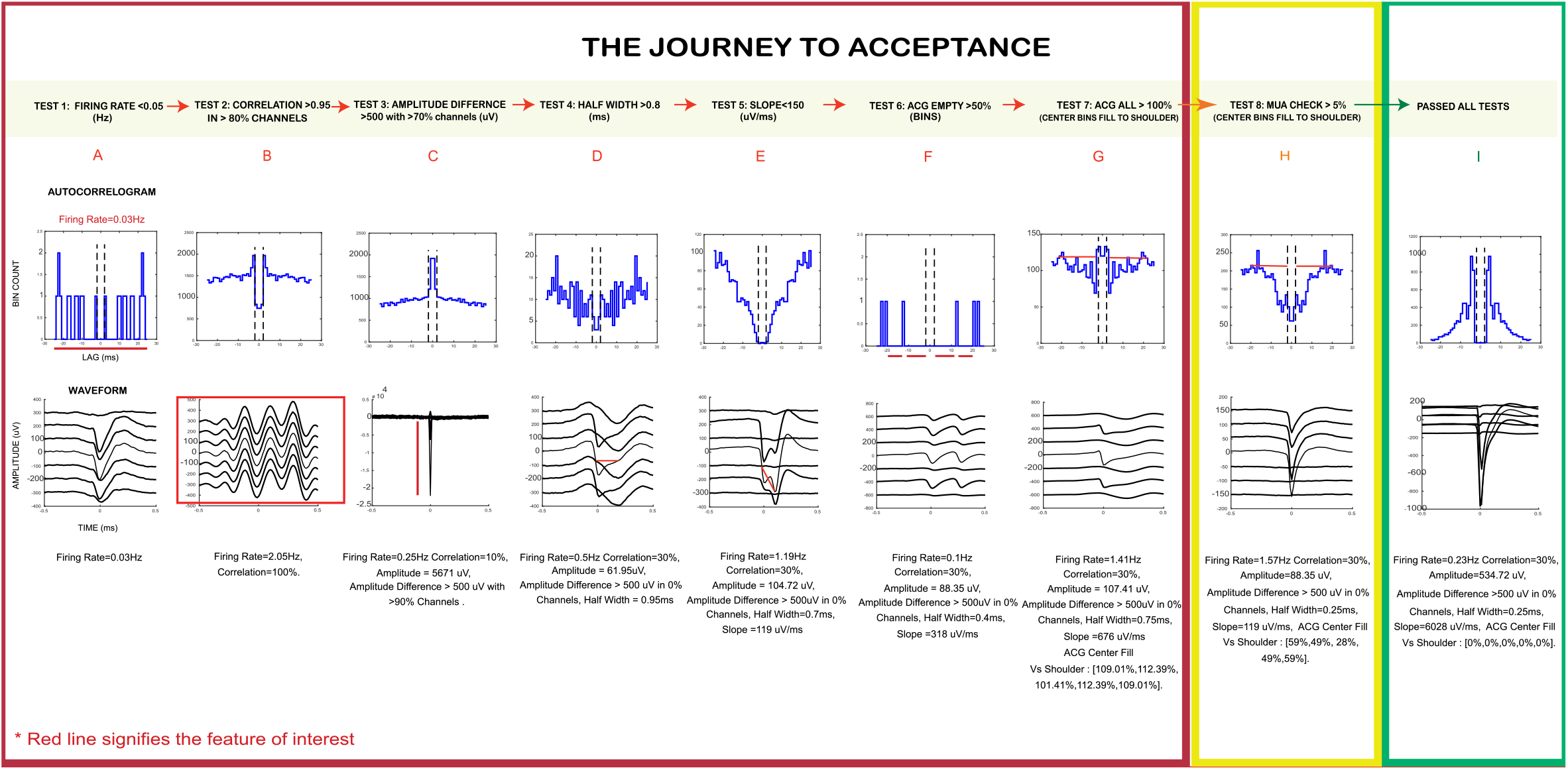
The Journey to Acceptance

**Figure 3.**
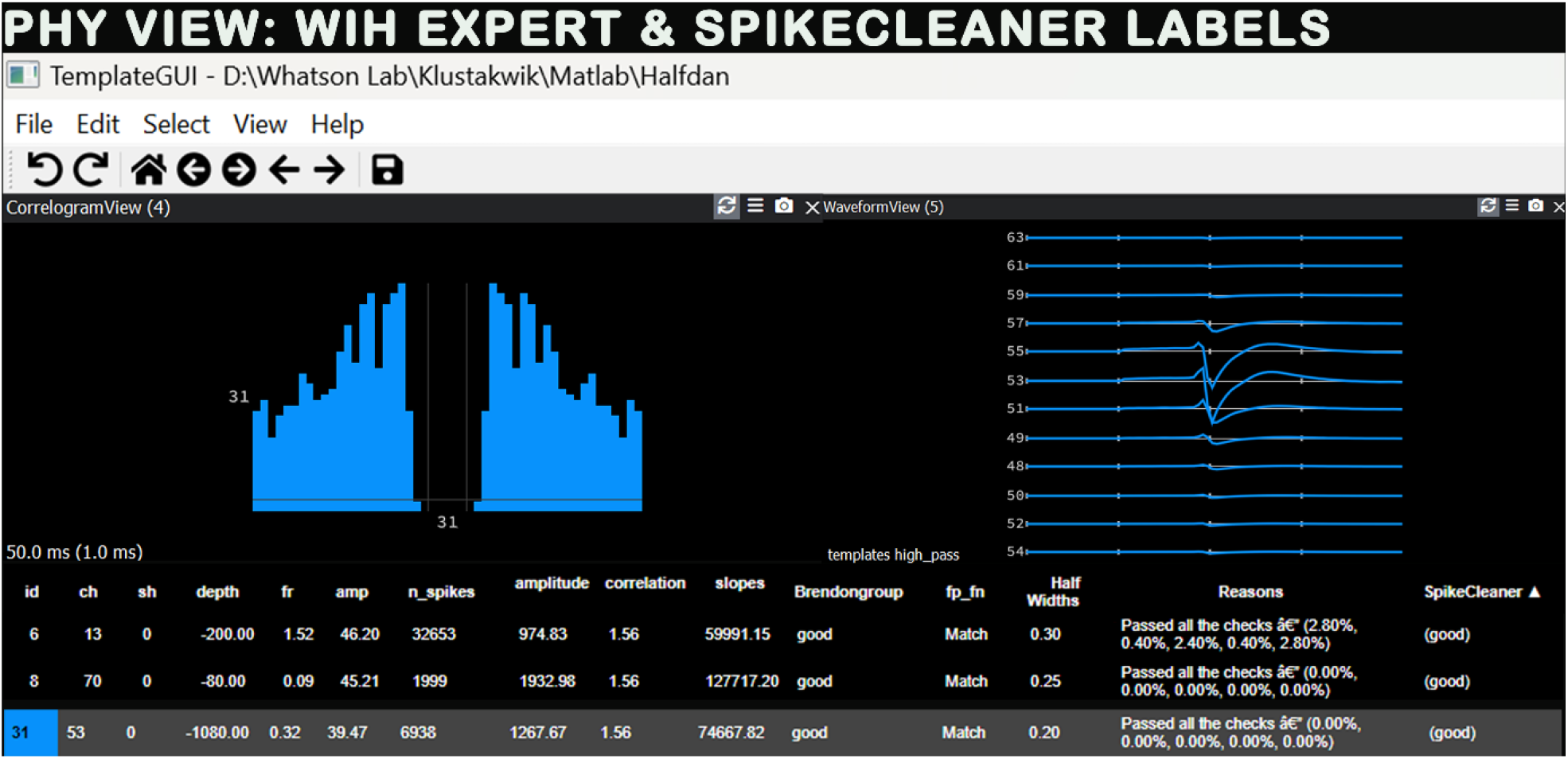
Visualization of SpikeCleaner Labels in PHY

**Figure 4.**
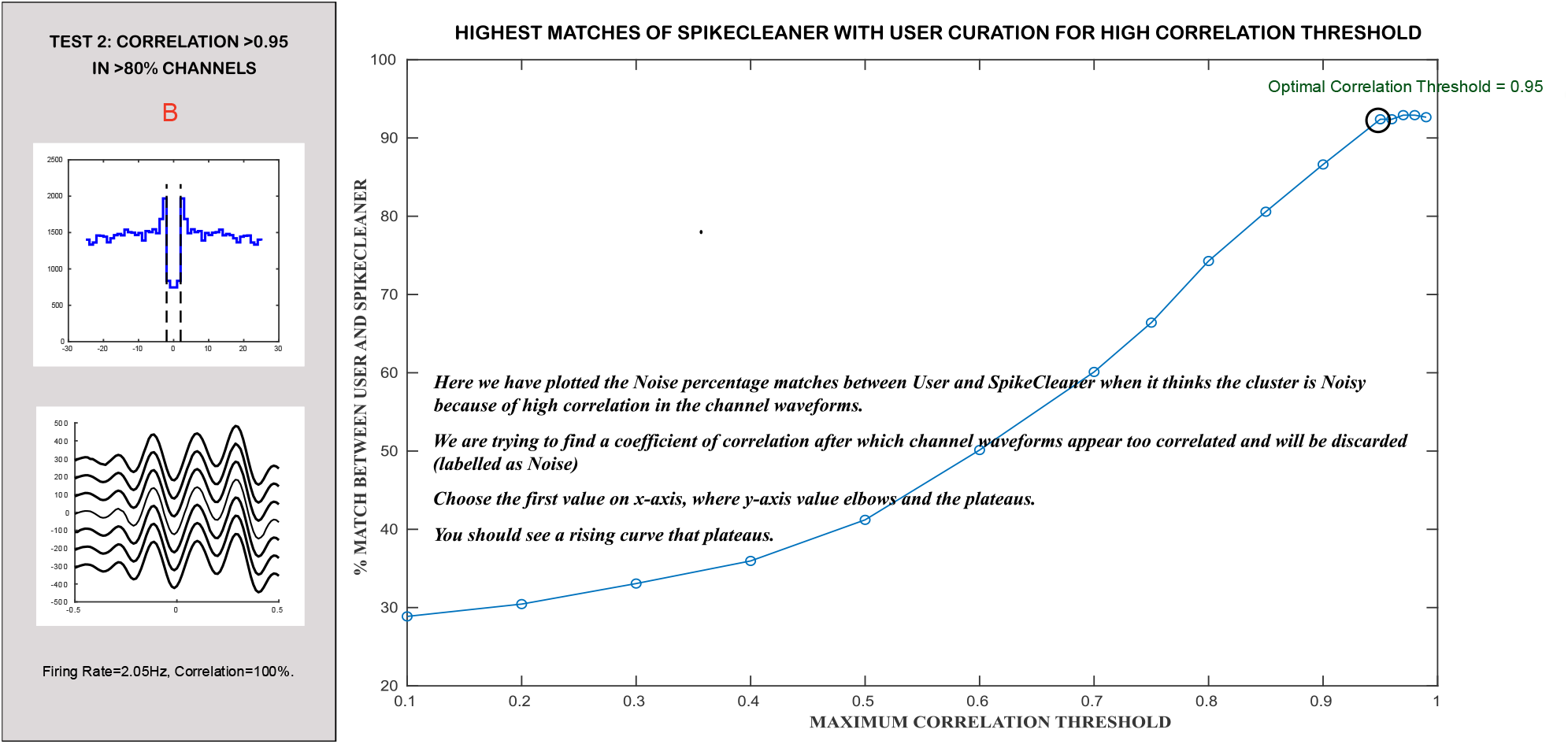
Determination of Correlation Threshold

**Figure 5.**
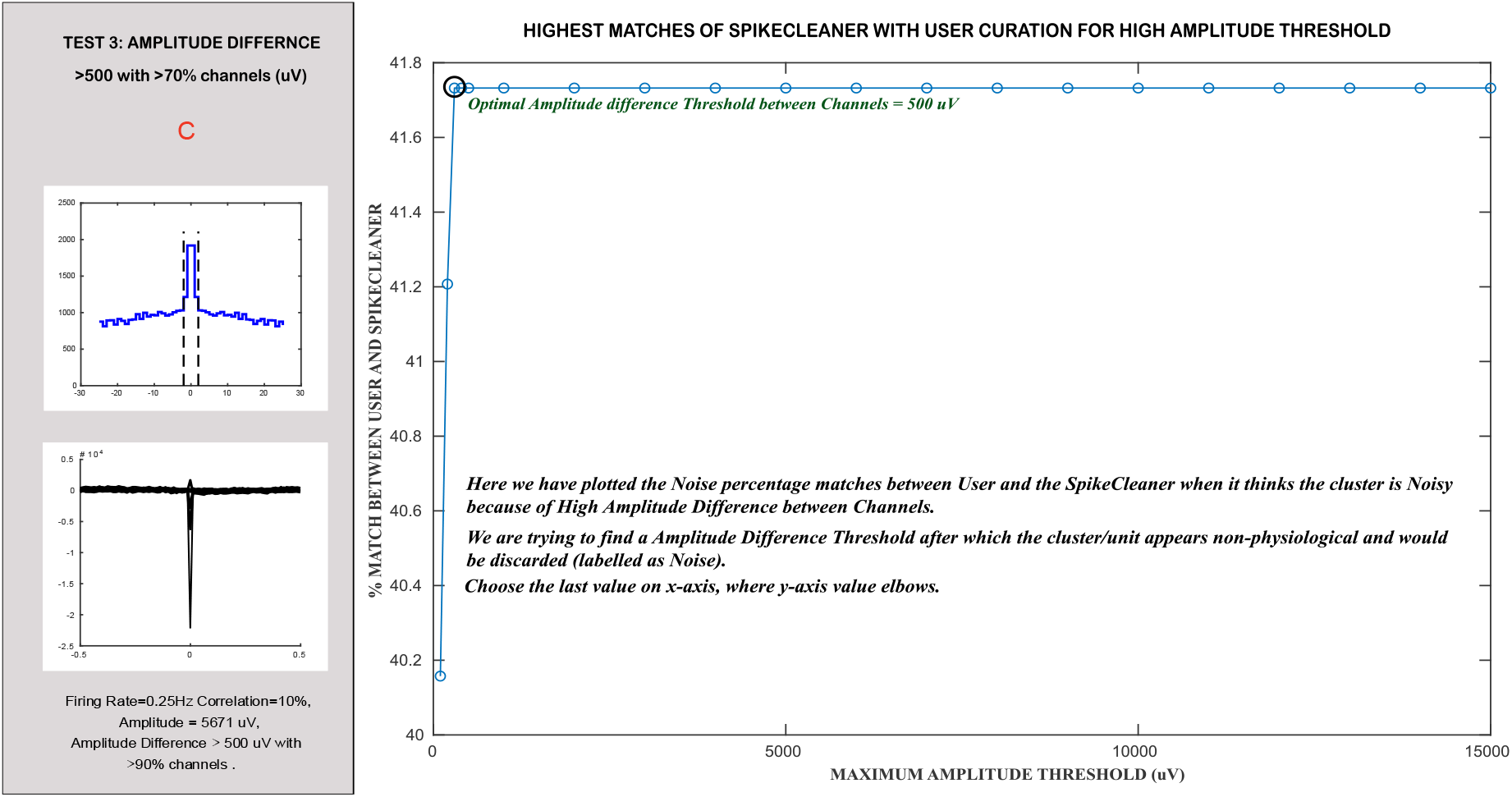
Determination of Amplitude Threshold

**Figure 6.**
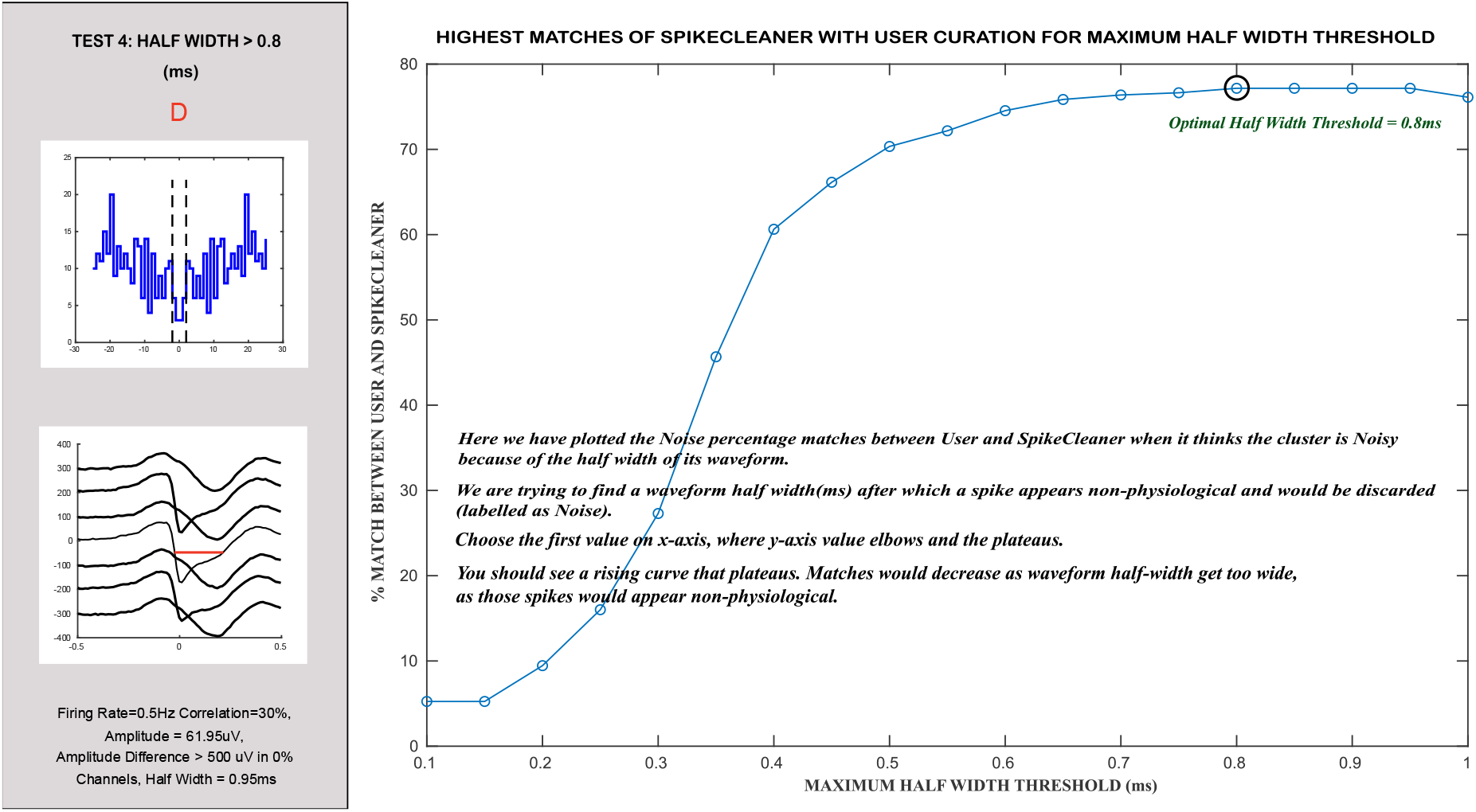
Determination of Half-Width Threshold

#### 2. Correlation Analysis

As illustrated in **Figure 2-B**, SpikeCleaner, performs a correlation screen between the maximum amplitude waveform and channels with less than 150 microns distance to it (closest in proximity), to eliminate clusters unlikely to represent biologically meaningful units. The electrode closest to the neuron would record the biggest signal and as electrode distance increases, signal amplitude decreases by the factor of 1/r^2^, ‘r’ being the distance between electrode and the firing neuron. Hence, physiologically accurate units have the largest spike at the channel closest to the neuron and then decreasing amplitude spikes in nearby channels. Having all channels at the same amplitude means there are some non-physiological aspects to the spike or its purely electrical noise.^**10**^

The correlation between maximum amplitude channel and all other channels is obtained from **dz_filterWaveform(). dz_Curate()** calls **dz_analyzeCorrelation()** from within the pipeline according to the order chosen by the user, it takes the **nLocalChannels**, a variable that user specifies and makes a subset of channels based on proximity to the position of the maximum amplitude channel. We obtain positions from **channel_positions.npy** (Output from Kilosort ^**1**^). It uses the **correlationthreshold** to analyse the percentage of high correlation channels. If a cluster has more than 80% of channels in selected channel range having higher correlation than the **correlationthreshold** to the maximum amplitude channel, it is labelled as Noise. Otherwise, the cluster progresses to the next step in the user chosen pipeline.

Threshold values for this criterion can be tuned for specific datasets using the threshold analysis scripts provided in **/SpikeCleaner/FindThreshold/findCorrelationThreshold**. For this user needs to first curate the dataset in PHY.

~~~
*
maxDistanceUm = 150; % use channels within 150 microns
%For every loop run:
if ∼good(ix)
  continue;
end
if isempty(corrValues)
  continue;
end
bestPos = pos(thisBestChannel,:);
d = sqrt(sum((pos - bestPos).^2,2));
 %using 150 microns
 % channels within 150 um, including best channel
 chRange = find(d <= maxDistanceUm);
 % get local correlations
 localCorrValues = corrValues(chRange);
 % remove self-channel
 localCorrValues(chRange == thisBestChannel) = [];
 totalChannels = length(localCorrValues);
 numHighCorr = sum(localCorrValues > correlationthreshold);
 percentHighCorr = (numHighCorr / totalChannels) * 100;
 if (percentHighCorr>80)
  noiseReasons{ix} = ‘Noise’;
  good(ix)=false;
else
  noiseReasons{ix} = ‘good’;
  good(ix)=true;
end
*
~~~

#### 3. Amplitude Analysis

As illustrated in **Figure 2-C**, SpikeCleaner, performs an amplitude screen between the maximum amplitude waveform and channels which are less than 150 microns to it (closest in proximity), to eliminate clusters unlikely to represent biologically meaningful units. We obtain positions from **channel_positions.npy** (Output from Kilosort ^**1**^). It consists of two checks, 1. The minimum amplitude channel is <50uV. ^**11**^ 2. The maximum amplitude channel shows an abnormally large spike that is not supported by corresponding activity on nearby channels that are flagged as likely artifacts and labelled as Noise.

**dz_Curate()** calls **dz_extractWfVariables** to extract the amplitude difference between the maximum amplitude channels and selected range of close by channels which is returned back as **chRangeAmp** structure for all the units. Then **dz_Curate()** calls **dz_analyzeAmplitude()** from within the pipeline according to the order chosen by the user, which checks if amplitude differences are higher than **maxAmp** threshold for more than 70% of channels in the selected range and labels the cluster as Noise. Otherwise, the cluster progresses to the next step in the user chosen pipeline.

Threshold values for this criterion can be tuned for specific datasets using the threshold analysis scripts provided in **/SpikeCleaner/FindThreshold/findAmplitudeThreshold**. For this user needs to first curate the dataset in PHY.

~~~
*
%for each loop run
if ∼good(ix)
  continue;
end
thisChRangeAmp=chRangeAmp{ix}; %in uV
Diff=thisAmplitude-thisChRangeAmp;
Diff(Diff == 0) = [];
totalChannels = length(thisChRangeAmp);
numHighAmpDiff = sum(Diff > maxAmp);
percentHighAmpDiff = (numHighAmpDiff / totalChannels) * 100;
if (percentHighAmpDiff>70) || thisAmplitude < minAmp
  noiseReasons{ix} = ‘Noise’;
  good(ix) = false;
else
  noiseReasons{ix} = ‘good’;
  good(ix) = true;
end
*
~~~

#### 4. Half-width Analysis

As illustrated in **Figure 2-D**, SpikeCleaner performs a half-width screening to eliminate clusters unlikely to represent biologically meaningful units. Biologically, extracellular action potentials are generated by rapid sodium influx followed by potassium-mediated repolarization, producing a brief voltage transient that typically lasts less than ∼ (1-2) ms. Waveforms that are substantially wider than this expected duration are unlikely to reflect single-neuron spiking activity and instead may arise from overlapping spikes, movement artifacts, filtering distortions, or low-frequency noise. Therefore, clusters with excessively large spike half-widths are excluded as non-physiological signals. ^**12**^

**dz_Curate()** calls **dz_extractWfVariables** to extract the half-widths of maximum amplitude waveform which returns a **halfwidths** structure for all the units. Then **dz_Curate()** calls **dz_analyzeHalfwidth()** from within the pipeline according to the order chosen by the user, which compares the half-width to **maxHW** threshold, and if the half-width is greater than the threshold, then the cluster is labelled as Noise. Otherwise, the cluster progresses to the next step in the user chosen pipeline.

Threshold values for this criterion can be tuned for specific datasets using the threshold analysis scripts provided in **/SpikeCleaner/FindThreshold/findHalfWidthThreshold**. For this user needs to first curate the dataset in PHY.

~~~
*
% for every loop run
thisHalfWidth=halfwidths{ix}; %in ms
if ∼good(ix)
  continue;
end
if thisHalfWidth<=maxHW
  noiseReasons{ix} = ‘good’;
  good(ix)=true;
else
  noiseReasons{ix} = ‘Noise’;
  good(ix)=false;
end
*
~~~

#### 5. Slope Analysis

As illustrated in **Figure 2-E**, SpikeCleaner performs a slope screening to eliminate clusters unlikely to represent biologically meaningful units. Biologically, extracellular action potentials are generated by rapid ionic currents, primarily fast sodium influx followed by potassium-mediated repolarization, resulting in steep rising and falling phases of the waveform. Waveforms with unusually shallow slopes reflect slower voltage changes that are inconsistent with the rapid dynamics of neuronal spiking and are more likely to arise from movement artifacts, low-frequency noise, or overlapping signals from multiple sources. Therefore, clusters with slopes below a physiological threshold are excluded as non-neuronal or unreliable signals.^**13**^

**dz_Curate()** calls **dz_extractWfVariables** to extract the slopes of maximum amplitude waveforms which return **slopes** structure for all the units. Then **dz_Curate()** calls **dz_analyzeSlope()** from within the pipeline according to the order chosen by the user, which labels clusters failing to meet the **minSlope** threshold as Noise. Otherwise, the cluster progresses to the next step in the user chosen pipeline.

Threshold values for this criterion can be tuned for specific datasets using the threshold analysis scripts provided in **/SpikeCleaner/FindThreshold/findSlopeThreshold**. For this user needs to first curate the dataset in PHY.

~~~
*
%for every loop run
if ∼good(ix)
  continue;
end
thisSlope=slopes{ix};%uV/ms
if thisSlope>=minSlope
  noiseReasons{ix} = ‘good’;
  good(ix)=true;
else
  noiseReasons{ix} = ‘Noise’;
  good(ix)=false;
end
*
~~~

#### 6. Empty ACG Analysis

As illustrated in **Figure 2-F**, SpikeCleaner performs an ACG screening to eliminate clusters unlikely to represent biologically meaningful units. A valid neuronal unit is expected to exhibit structured spike timing patterns in its ACG, reflecting refractory period suppression, near zero lag, and consistent firing relationships over short time intervals. Clusters whose ACG contains mostly zero or near-zero values lack sufficient temporal structure and typically arise from extremely low firing rates, noise contamination, or unstable spike detection. Such clusters are therefore flagged as unreliable and excluded from further analysis.

**dz_Curate()** calls **dz_extractAcgVariables** to extract the proportions of center bins with respect to the shoulder bins and if the ACGs are more than 50% empty for all the units. Then **dz_Curate()** calls **dz_acgEmpty()** from within the pipeline according to the order chosen by the user, if the value of **isEmpty** is true for a cluster it’s labelled as Noise, otherwise the cluster progresses to the next step in the user chosen pipeline.

~~~
*
%for every loop run
if ∼good(ix)
  continue;
else
thisEmpty=isEmpty(ix);
thisProportion=Proportions{ix};
if thisEmpty==true
  good(ix)=false;
  noiseReasons{ix} = ‘Noise’;
end
*
~~~

#### 7. ACG-all Analysis

As illustrated in **Figure 2-G**, SpikeCleaner performs an ACG screening to eliminate clusters unlikely to represent biologically meaningful units. This step evaluates the ratio between spike counts in the central ACG bins (±2 ms around zero lag) and the shoulder region of the ACG. These center-to-shoulder proportions indicate how strongly the refractory period is preserved. Clusters with central bins that are comparable to or larger than the shoulder bins typically reflect refractory violations and are therefore likely to represent multi-unit activity (MUA) or poorly isolated noise clusters, while biologically meaningful single units are expected to show reduced spike counts near zero lag due to refractory period suppression. Threshold values for this criterion can be tuned for specific datasets using the threshold analysis scripts provided in **/SpikeCleaner/FindThreshold/findACGallThreshold**.

**dz_Curate()** calls **dz_extractAcgVariables** to extract the proportions of center bins with respect to the shoulder bins and if the ACGs are more than 50% empty for all the units. Then **dz_Curate()** calls **dz_acgAllCheck()** from within the pipeline according to the order chosen by the user, if the value of **Proportions** are greater than **acgallthreshold** then the cluster is labelled as Noise/MUA depending on the **acgalllabel** value that the user obtained from running **findACGallThreshold()**, otherwise the cluster progresses to the next step in the user chosen pipeline.

Threshold values for this criterion can be tuned for specific datasets using the threshold analysis scripts provided in **/SpikeCleaner/FindThreshold/findACGallThreshold**. For this, users need to first curate the dataset in PHY.

~~~
*
%for every loop run
if ∼good(ix)
  continue;
else
thisProportion=Proportions{ix};
if all(thisProportion>acgallthreshold)
  good(ix)=false;
  noiseReasons{ix} = acgalllabel;
else
  good(ix)=true;
  noiseReasons{ix} = ‘good’;
end
*
~~~

#### 8. Multi-Unit Activity (MUA) Analysis

As illustrated in **Figure 2-H**, SpikeCleaner performs an ACG screening to identify and separate multi-unit activity (MUA) clusters from well-isolated single units. Unlike single units, which exhibit a clear refractory period suppression near zero lag, MUA clusters typically show elevated spike counts in the central ACG bins due to contributions from multiple nearby neurons firing independently. By comparing the structure of the ACG around the zero-lag region to the shoulder bins, SpikeCleaner further distinguishes mixed quality clusters and classifies such clusters as MUA rather than single-unit activity, improving the reliability of downstream analyses.

**dz_Curate()** calls **dz_extractAcgVariables** to extract the proportions of center bins with respect to the shoulder bins and if the ACGs are more than 50% empty for all the units. The proportion is the ratio between the values of (±2ms) center bins and the shoulder bin. The shoulder bin is calculated as the average of most prominent bins after the center bins. Then **dz_Curate()** calls **dz_muaCheck ()** from within the pipeline according to the order chosen by the user, which applies either the **strict** or **lenient** rules for the MUA check depending on the **acgEvaluationMode** chosen. The default is set to ‘**strict’**.

In the **strict acgEvaluationMode**, all proportion values are expected to be less than 5%, that is, only 5% fill is allowed in the center bins with respect to the shoulder bins. In the **lenient acgEvaluationMode, (**±2ms) th bin’s **proportion** values need to be lower than 30% and (±1ms) bin’s and 0th bin’s **proportion** values need to be lower than 20%. If the ACG fails, this criterion it will not pass to be a **single unit (SU)/good** and will be labelled as MUA.

~~~
*
%for every loop run
switch acgEvaluationMode
 case ‘lenient’
  %±2 ms bins must be < 30
  if thisProportion(1) > 0.3 || thisProportion(6) > 0.3
   flag = true;
  %Allow 0th and ±1ms bins to be < 30% if outer bins are as low as 30
  elseif (thisProportion(1) < 0.3 && thisProportion(6) < 0.3) && all(thisProportion(2:5) < 0.2)
   flag = false;
  else
   flag = true;
  end
case ‘strict’
  if any((thisProportion*100)>0.05) % 5%
   flag = true;
  else
   flag=false;
  end
end
if flag==true
  good(ix)=false;
  noiseReasons{ix} = ‘MUA ‘;
  continue;
end
*
~~~

#### SpikeCleaner Outputs

SpikeCleaner outputs in the **SpikeCleaner/**, curation labels in **cluster_SpikeCleaner.tsv**, detailed curation reasons in **cluster_Spikereasons.tsv**, along with this, it also outputs all computed feature metrics (e.g., ACG-based measures, waveform properties, firing rate, correlation features in tsv formats in the **SpikeCleaner** folder. These outputs allow users to seamlessly load results into Phy for review while providing transparent reasoning and quantitative metrics behind each label. The goal is to assist users in analyzing their data more efficiently, reproducibly, and in a standardized manner across datasets and curators. All the parameters, thresholds and pipeline flow from the current run are saved in **animalname_SpikeCleaner_settings.json** in **SpikeCleaner/** which can be edited or one can make multiple versions of it and change the input file based on the experiment, for this the user would need to update the file name in **dz_classifyAllUnits()**

SpikeCleaner generates summary figures that are saved in the **SpikeCleaner/** output directory to help users evaluate the quality of their recordings based on the number of SU/Good Units, MUAs and Noisy units obtained and assess how each step in the pipeline contributes to cluster refinement. These visualizations show how many clusters are removed at each stage, allowing users to identify which criteria are most influential for their dataset. For example, if a large proportion of clusters are discarded during the correlation check, that step can be placed earlier in the pipeline to avoid unnecessary computation in later stages. This makes the workflow more efficient and allows users to optimize the sequence of quality-control steps for their specific recordings (see results).

#### Visualization of Results in PHY

From a user interface perspective, we have designed SpikeCleaner output to be read by Phy. This reduces the need for extra software, weaving SpikeCleaner into the common Kilosort ^**1**^-Phy ^**6**^ pipeline used by many labs. Furthermore, we integrate with Phy in two specific ways. First, SpikeCleaner only <u>suggests</u> labels for users to later confirm, rather than automatically labeling, thereby leaving crucial human judgment possible. Second, via modification of the Phy-readable .csv file for a given recording, the SpikeCleaner reasons for a given decision are displayed for the user (see the second column from right in Figure 7). This is a “suggestion-only” feature, it creates a new header for SpikeCleaner labels and does not interfere with the user labels. Once a user has calibrated the thresholds and has established trust, they can choose to run SpikeCleaner without manual curation.

**Figure 7.**
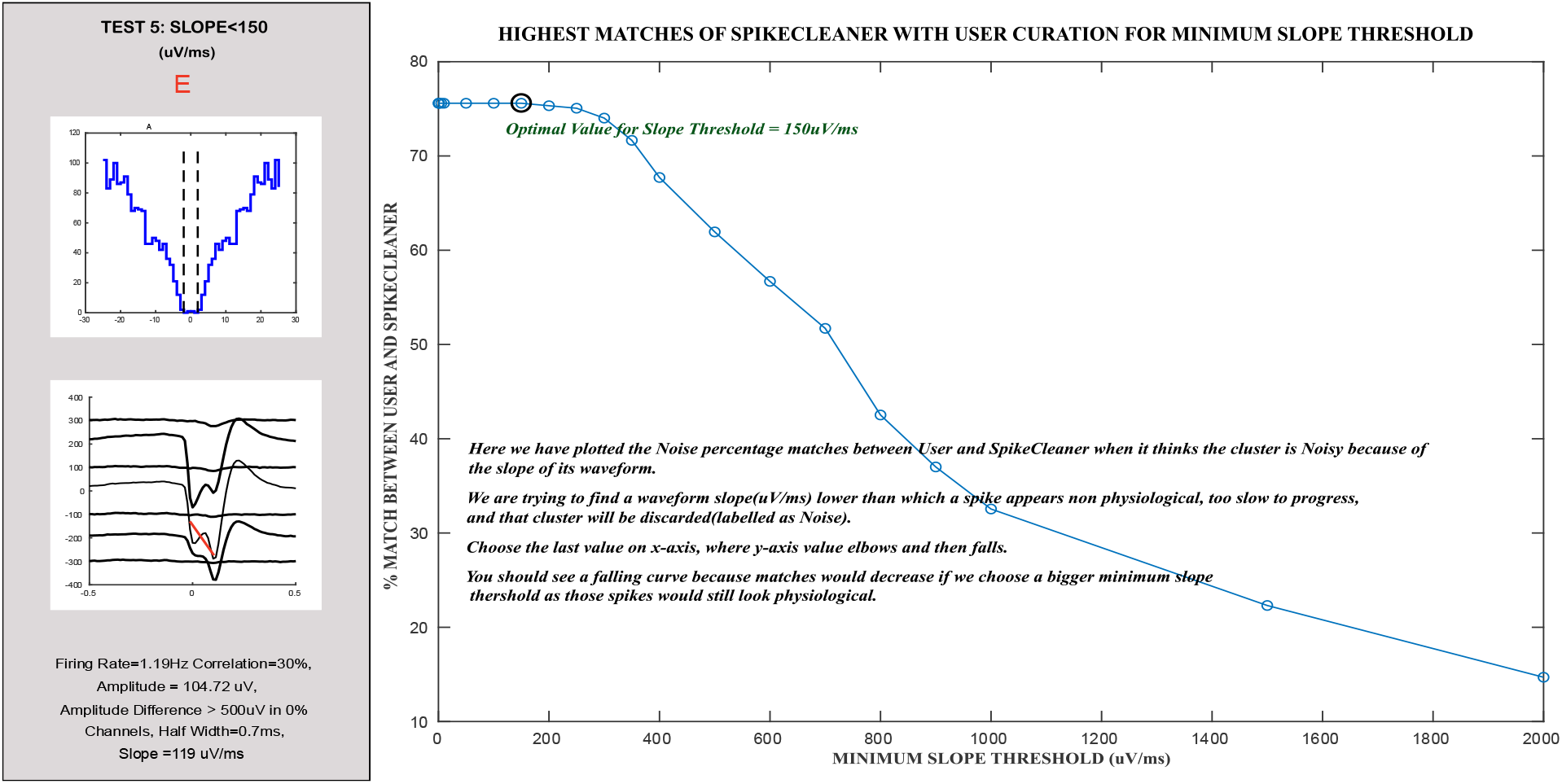
Determination of Slope Threshold

**Figure 8.**
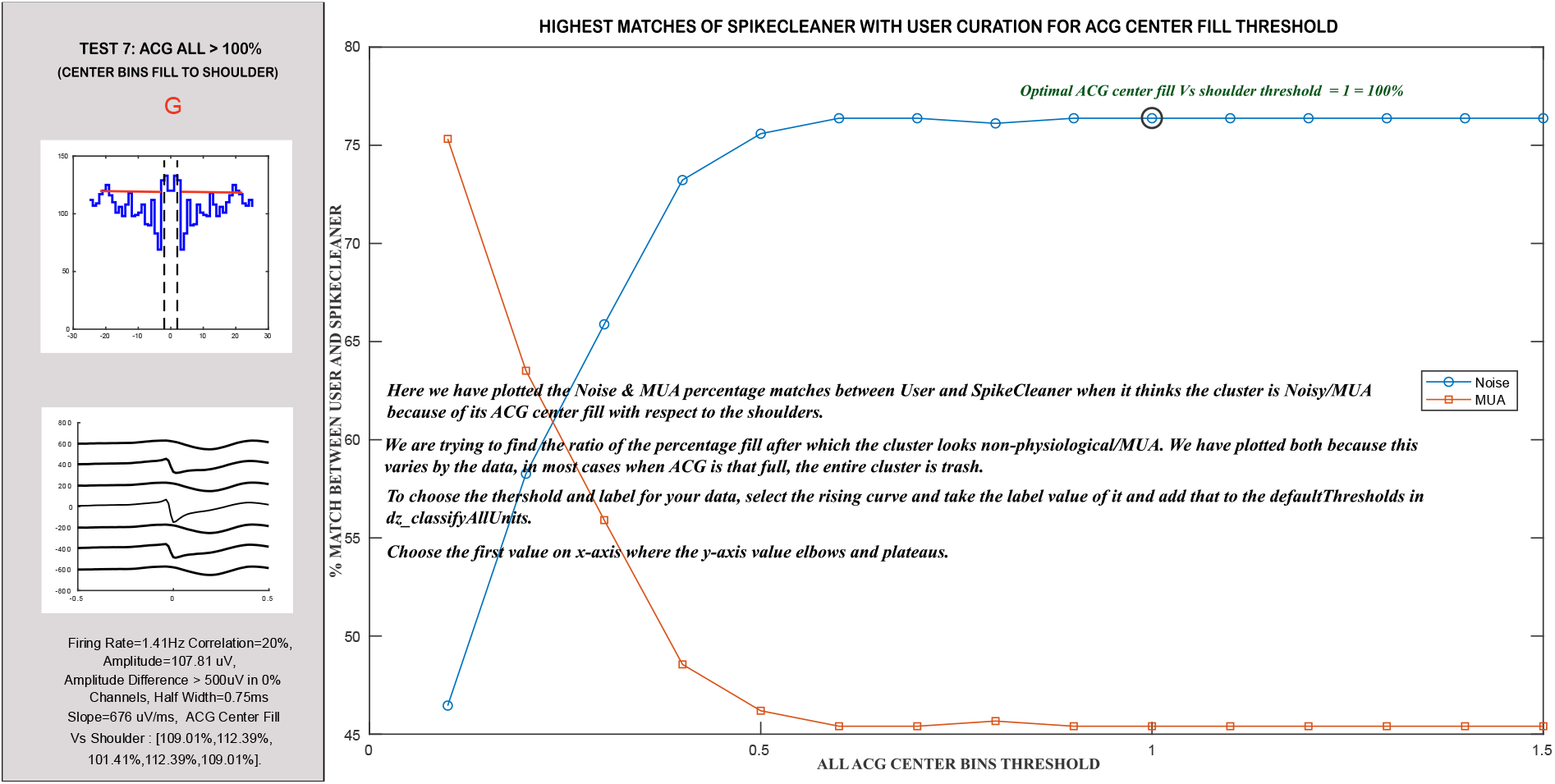
Determination of Threshold for all ACG Center Bins

A key strength of SpikeCleaner is that all decision thresholds, including those for amplitude difference, half-width, slope, spatial correlation, and ACG-based refractory violations, are fully adjustable, allowing users to tune the algorithm to match their own curation style or the characteristics of a specific dataset. As depicted in Figure 7, SpikeCleaner outputs detailed per-cluster metrics and generates Phy-readable labels; users can iteratively modify thresholds.

Furthermore, if users save fully human-scored expert classifications of noise/mua/good from Phy for a given recording, they can immediately compare the resulting classifications to their own manual curation, or to reference expert curators. The user needs to run **SpikeCleaner/AccuracyMetrics/** codes inside the **SpikeCleaner/** in the data folder. The code modules: **dz_goodvsRest()**,**dz_allCats()**,**dz_NeuronalvsNN()**, asks user to enter their name and renames the user curation file **cluster_group.tsv** to **cluster_group_username.tsv and** changes the column header inside from **group** to **usergroup** to ensure clear differentiation of user labels and SpikeCleaner labels in PHY. It outputs **goodvsRest_username.tsv, allcat_username.tsv, NvsNN_username.tsv** in **SpikeCleaner/** which are Match/No Match metrics for each of the categories. These outputs are imported in PHY and a user can easily sort the clusters based on them. It also outputs a table for accuracy statistics between user and SpikeCleaner for each of the categories in **SpikeCleaner/Accuracy/**

### Obtaining Quality Check Thresholds Customized to User’s Data and Determination of Default Thresholds

To determine optimal feature thresholds, we evaluated the performance of SpikeCleaner across a range of parameter values by comparing its classifications to gold-standard manual curation from two expert annotators. We measured the algorithm’s agreement with expert labels across a range of possible thresholds. Specifically, we compared the cluster_Spikereasons.tsv output file, which records the criteria responsible for labeling clusters as Noise or MUA, against the corresponding user-curated labels. This comparison allowed us to systematically identify threshold values that maximize agreement with experts scoring across all quality-control steps.

#### 1. Correlation Threshold

This plot shows the percentage agreement between SpikeCleaner and expert manual curation as a function of the maximum waveform correlation threshold used to classify clusters as noise. As the threshold increases, agreement improves and reaches a plateau near 0.95, indicating the point beyond which additional increases provide minimal benefit. Based on this elbow– plateau behavior, a correlation threshold of 0.95 was selected as the optimal value for identifying clusters with excessively correlated channel waveforms that likely reflect artifacts rather than biologically meaningful units.

#### 2. Amplitude Threshold

Agreement between SpikeCleaner and expert curation increases rapidly at low amplitude-difference thresholds and plateaus near 500 µV, indicating that clusters with extremely large inter-channel amplitude differences are reliably identified as non-physiological. Thresholds beyond this value provide minimal additional improvement, supporting 500 µV as an appropriate cutoff for artifact rejection.

#### 2. Half-Width Threshold

This plot shows the percentage agreement between SpikeCleaner and expert manual curation as a function of the waveform half-width threshold used to classify clusters as noise. Agreement increases as the threshold approaches physiologically realistic spike durations and reaches a plateau near 0.8 ms, indicating the point beyond which wider waveforms are increasingly likely to reflect non-physiological signals such as overlapping spikes, movement artifacts, or low-frequency contamination. Based on this elbow–plateau behavior, a half-width threshold of 0.8 ms was selected as the optimal cutoff for rejecting clusters with unusually broad spike waveforms.

#### 3. Slope Threshold

Agreement remains relatively stable for low slope thresholds, indicating that physiologically realistic spikes are retained despite increasingly strict filtering. However, beyond approximately 150 µV/ms, agreement begins to decline sharply, suggesting that valid neuronal waveforms are increasingly misclassified as noise. Waveforms with slopes below this value typically exhibit unusually slow rise and fall dynamics that are inconsistent with action potentials and may instead reflect overlapping spikes, movement artifacts, or low-frequency contamination. Based on this elbow-like transition in the agreement curve, a minimum slope threshold of 150 µV/ms was selected as the optimal cutoff for identifying non-physiological waveforms while maximizing agreement with expert curation.

#### 4. ACGall Threshold

This plot shows the agreement between SpikeCleaner and expert manual curation as a function of the ratio between spike counts in the central ACG bins (±2 ms around zero lag) and the shoulder region of the autocorrelogram. Single-unit activity is expected to exhibit reduced spike counts near zero lag due to refractory period suppression, whereas clusters with central bins comparable to or exceeding shoulder bins typically indicate refractory period violations consistent with multi-unit activity (MUA) or noise. Agreement with expert labeling increases as the center-to-shoulder ratio approaches 1 and then plateaus, that is the center bins at or above 100% of the shoulder bin, indicating that clusters with fully filled central ACG bins are unlikely to represent well-isolated single units. Depending on the dataset, elevated center-bin filling may correspond to either MUA or noise; therefore, users should select the label associated with the rising portion of the curve (Noise or MUA) and update the **acgalllabel** parameter while calling **dz_classifyAllUnits()** accordingly to match their recording characteristics.

The user needs to first manually curate the recording and then run code modules in **SpikeCleaner/FindThresholds/ i**n the **code folder** and output plots are saved in **SpikeCleaner/Correlation, SpikeCleaner/Amplitude, SpikeCleaner/Slope, SpikeCleaner/HalfWidth/, SpikeCleaner/ACGall in the data folder**.

These obtained thresholds can be used to call **dz_classifyAllUnits():**

For this the user would need to add/change the thresholds in **animalname_SpikeCleaner_settings.json** in SpikeCleaner/ or create a new json file and update the name in **dz_classifyAllUnits**.

This systematic threshold-sweeping approach ensures that our final cutoff is both physiologically grounded and empirically optimized for expert-level performance. For example, sweeping the half-width threshold revealed that 0.8 ms produced the closest agreement with expert-labeled biologically possible units (Good+MUA) (>75% match), demonstrating how threshold selection can be data-driven rather than arbitrary. In this way, SpikeCleaner serves not only as an automated curation tool but also as a calibration framework that allows laboratories to standardize, document, and reproduce expert curation criteria to their own specifications should they desire to fine-tune it.

The experts reviewed individual unit disagreements between SpikeCleaner and experts, and the conclusion was that disagreements were within the range of “reasonable disagreement” one might expect between individual human curators. There did not appear to remain systematic deviation between experts and the algorithm. Given the nature of the differences, we did not modify this further.

## RESULTS

SpikeCleaner follows a sequential, rule-based pipeline designed to progressively eliminate non-biological clusters and isolate high-quality single units. The algorithm first removes clusters with implausibly low firing rates, then extracts and filters average waveforms from the best channel of each remaining cluster. Waveform shape is reduced to a three-point extremum triplet, from which amplitude, half-width, and pre/post post-slopes are computed and thresholded to reject non-physiological or low-quality units. A spatial correlation check ensures that true neuronal waveforms exhibit localized spread rather than globally synchronized noise. Finally, an autocorrelogram-based assessment classifies units as Single Units, Multi-Unit Activity, or Noise by quantifying refractory period violations and central-bin filling. Together, these sequential evaluations form a standardized, transparent curation pipeline that emulates expert decision-making while maintaining reproducibility and speed.

### Recordings for validation

To test this pipeline, we have applied it to and tested against three animal recordings of 20 kHz, 16-bit, 128-channel continuous recording, spanning approximately 6 hours and producing ∼110 GB of raw data per session. Spike sorted by Kilosort ^**1**^. We recorded using 64-channel headstages (RHD2132 by Intan Technologies, Los Angeles, CA) connected to a USB interface board (RHD2000 by Intan Technologies, Los Angeles, CA) and sampled at 20 kHz. Datasets for this study were acquired from the linear silicon probes targeted at 24h continuous recordings with freely moving animals.

Three animals were used in this study. All procedures were performed in accordance with the guidelines of the University of Michigan Institutional Animal Care and Use Committee (IACUC). Male Nile grass rats (*Arvicanthis niloticus*) were obtained from the long-established breeding colony maintained at Michigan State University by Dr. Lily Yan.^**8**^ Animals arrived at 8– 12 weeks of age and were allowed to acclimate to the University of Michigan vivarium for at least three weeks before electrophysiological experiments.

Neural activity was recorded using linear 1×64 silicon probes (ASSY-325 H3, Cambridge NeuroTech, Cambridge, UK) mounted onto microdrive assemblies (Nano-Drive v1, Cambridge NeuroTech). Probes were stereotaxically implanted under isoflurane anesthesia using a Kopf arm (Kopf Instruments, Tujunga, CA, USA) into two target regions: the medial prefrontal cortex (mPFC; from bregma: A/P +0.48 cm, M/L +0.03 cm, D/V −0.32 cm) and dorsal hippocampus (dHC; from bregma: A/P +0.01 cm, M/L +0.20 cm, D/V −0.16 cm). Recordings were conducted after a week of recovery and were chronic, allowing multi-day monitoring of neural activity on freely moving animals. Each dataset consisted of a 20 kHz, 16-bit, 128-channel continuous recording, spanning approximately 6 hours and producing ∼110 GB of raw data per session. These large, high-density datasets provided a rigorous test of SpikeCleaner’s scalability and robustness and were used in benchmarking tests. All initial spike sorting was carried out with Kilosort 1. ^**1**^

### Comparison against manual curation

Manually curated by two experts for benchmarking. As illustrated in **Figure 9**, the algorithm achieved **expert-level performance** across all evaluation categories. For distinguishing **Single Units vs. Rest**, SpikeCleaner reached an accuracy of **97%** with an **F1 score of 92%**, indicating strong precision and recall in isolating high-quality single units. In the full three-class comparison (**SU vs. MUA vs. Noise**), the algorithm maintained similarly high performance, achieving **97% accuracy** and a **92% F1 score**. When collapsing classes into **Biological (SU+MUA) vs. Non-Biological (Noise)**, SpikeCleaner achieved an accuracy of **97%** and an **F1 score of 95%**, demonstrating robust discrimination between biological and non-biological clusters. These results show that SpikeCleaner closely replicates expert decision-making while providing a fast, standardized, and fully reproducible alternative to manual curation

**Figure 9.**
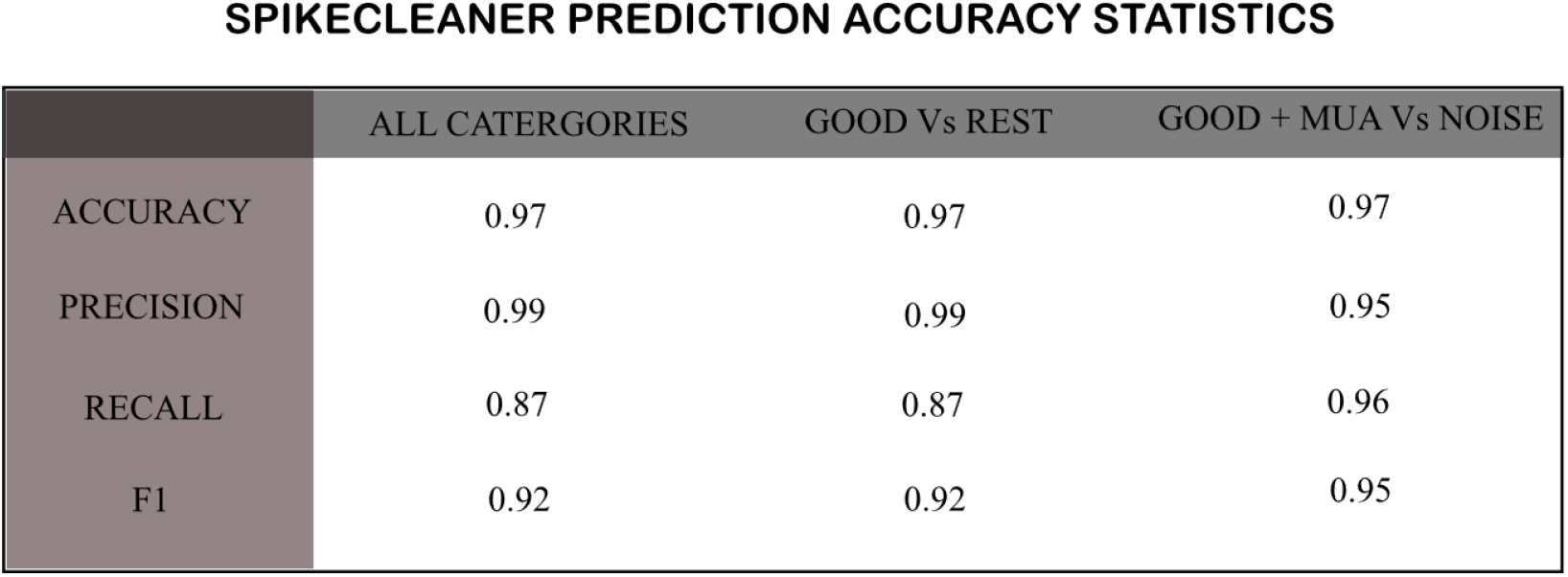
Prediction Accuracy Metrics

**Figure 10.**
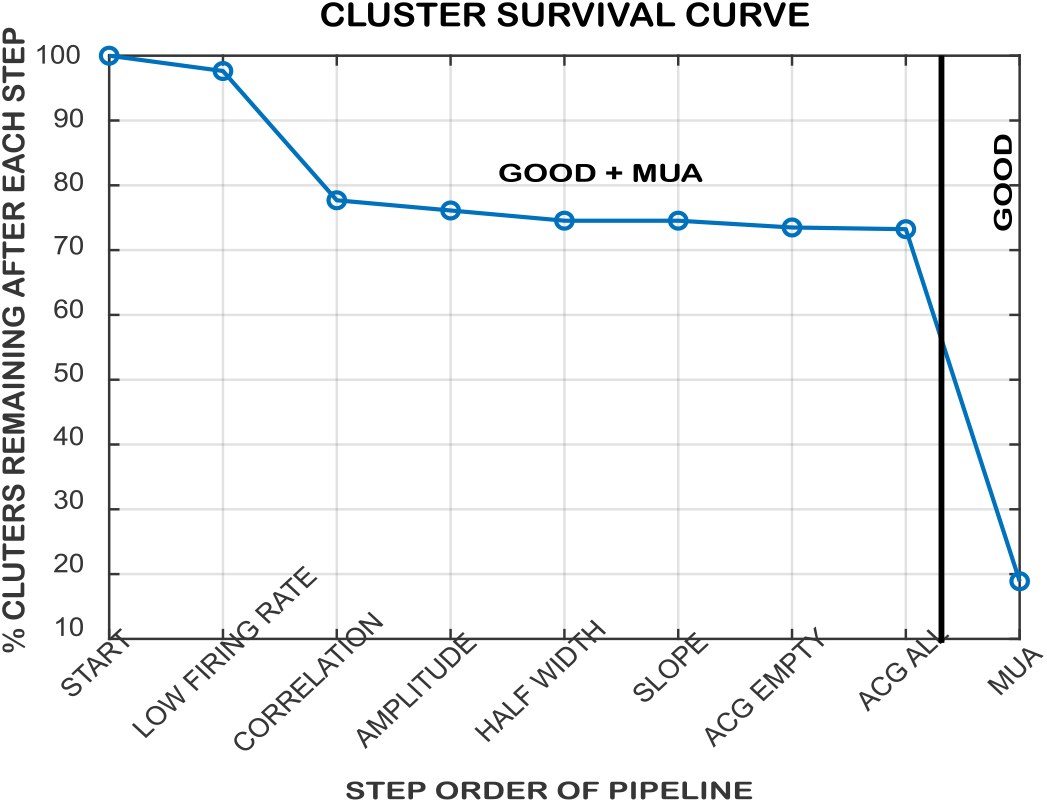
Cluster Survival Curve

**Figure 11.**
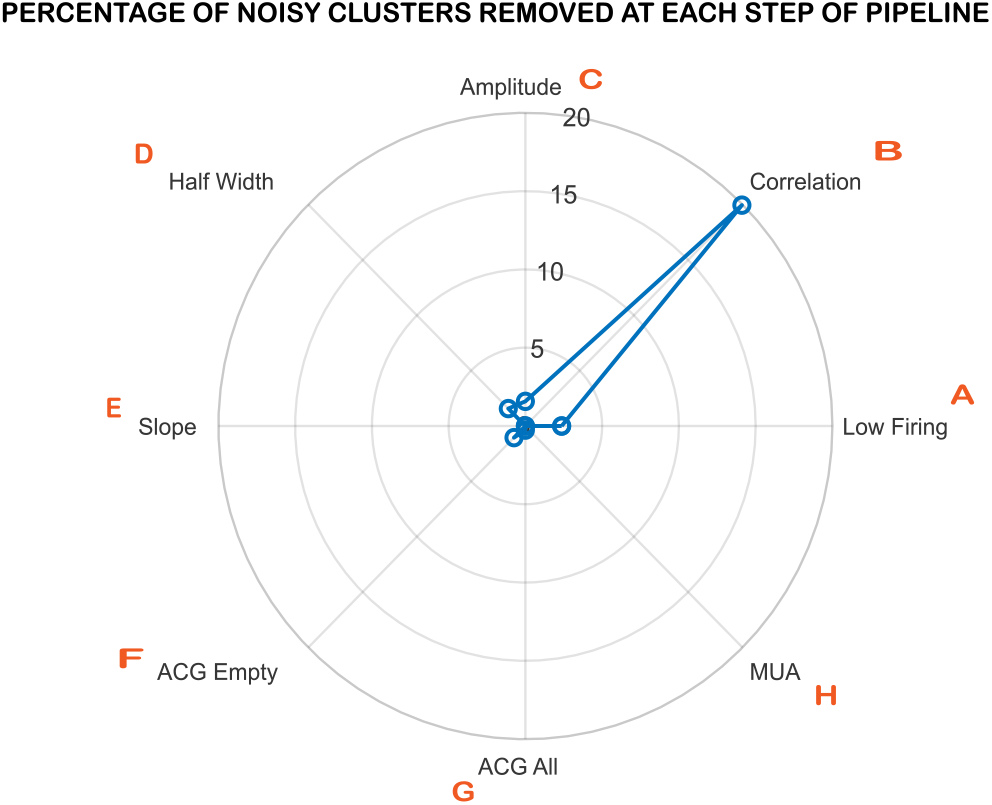
Percentage of Noisy Clusters Removed per Step

**Figure 12.**
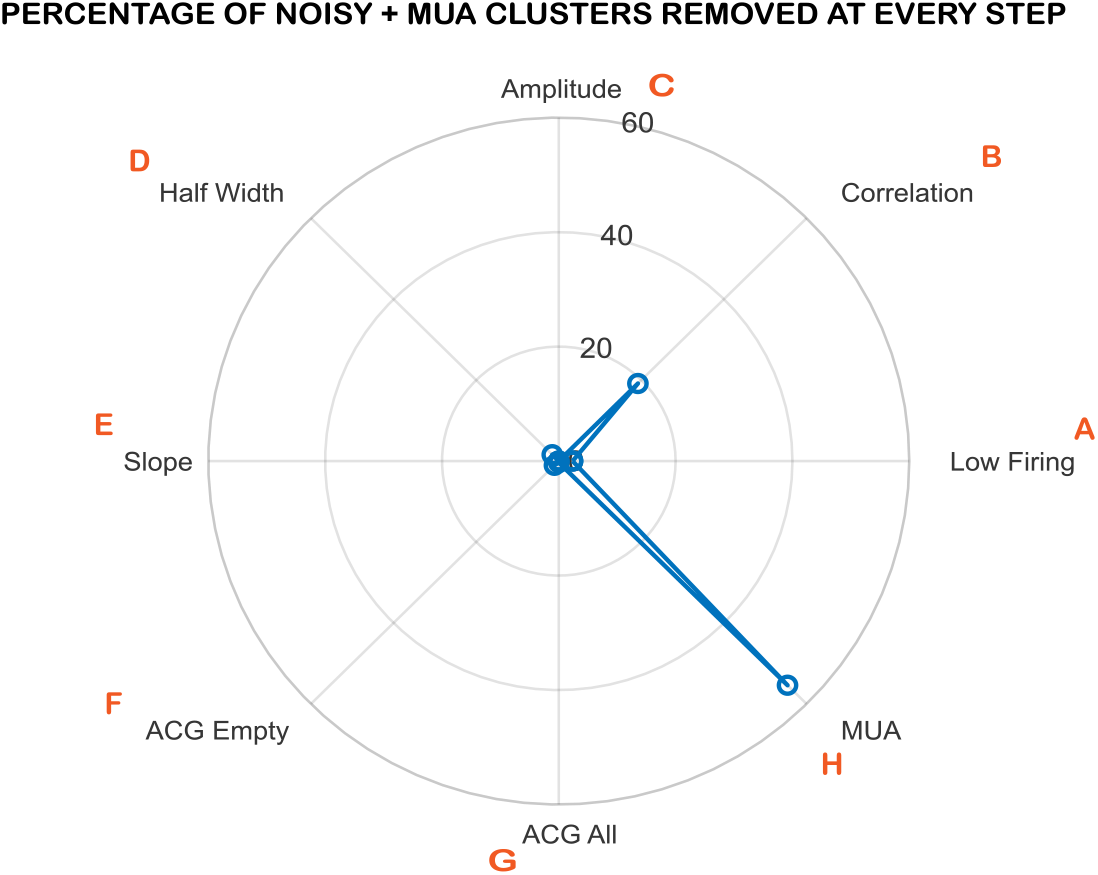
Percentage of Non–Single Unit Clusters Removed per Step

**Figure 13.**
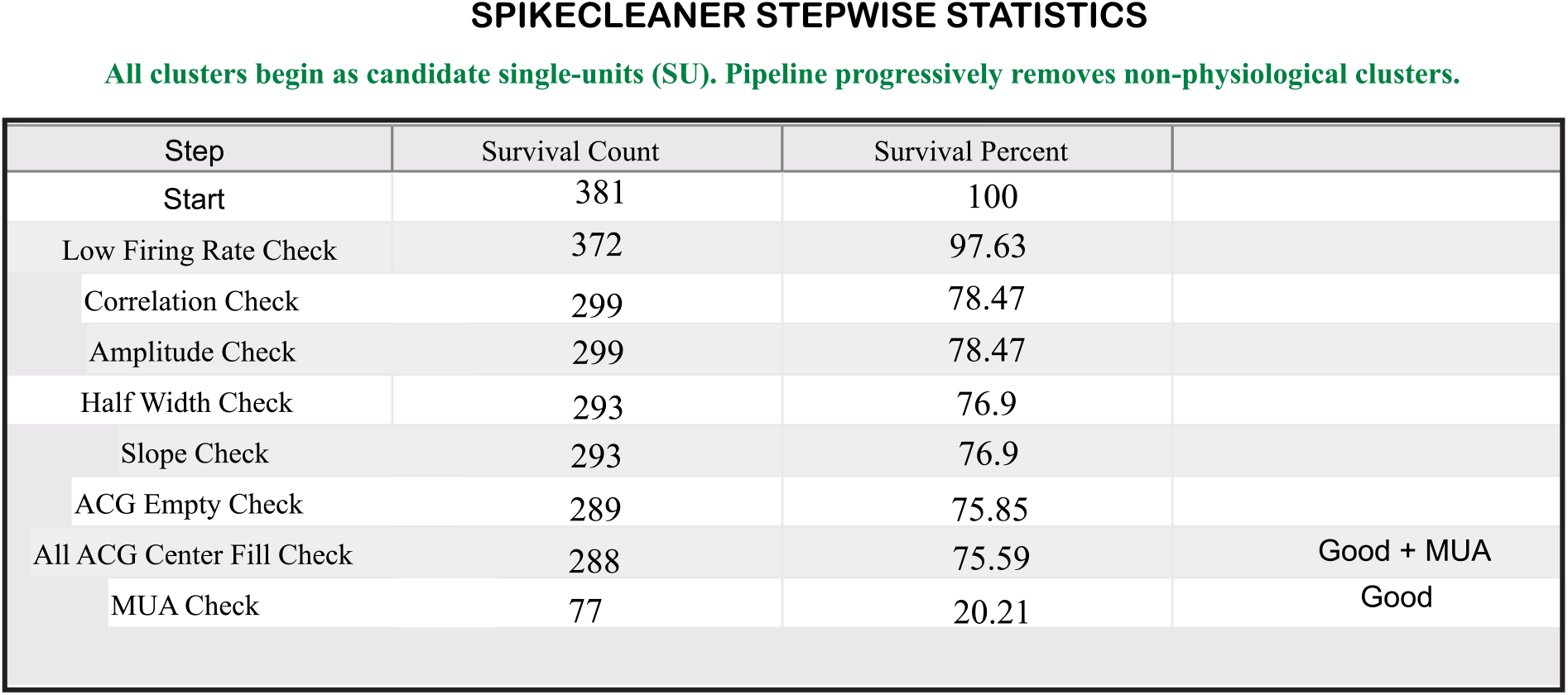
Stepwise Survival Statistics Table

### SpikeCleaner generates summary figures

This figure shows the percentage of clusters remaining after each step of the SpikeCleaner pipeline. All clusters initially enter the pipeline as candidate single units, and successive filtering steps progressively remove clusters identified as noise or non-physiological. The curve provides an overview of how each stage contributes to refining the dataset and illustrates the cumulative effect of quality-control criteria across the pipeline. The survival curve is generated based on the default pipeline order and is automatically updated if the sequence of pipeline steps is modified, allowing users to efficiently optimize the workflow for their specific recordings.

This radar plot shows the proportion of clusters removed at each individual pipeline step based on biological vs non-biological classification, that is, MUAs+Good/SUs. It highlights which screening criteria (e.g., amplitude, slope, correlation, ACG checks) contribute most strongly to eliminating noisy or unreliable clusters. This helps users understand which quality-control features are most influential for their dataset. The plot is generated based on the default pipeline order and is automatically updated if the sequence of pipeline steps is modified, allowing users to efficiently optimize the workflow for their specific recordings.

This radar plot specifically tracks removal of clusters that fail single-unit criteria but may still include multi-unit activity. It shows how each pipeline step improves isolation quality by separating well-isolated single units from partially overlapping neuronal activity. This provides insight into which steps most strongly refine single-unit classification. The plot is generated based on the default pipeline order and is automatically updated if the sequence of pipeline steps is modified, allowing users to efficiently optimize the workflow for their specific recordings.

This table summarizes the number and percentage of clusters remaining after each pipeline step. It reports:

- Total neuronal clusters (Good + MUA)
- Isolated single units (Good only)
- Percentages relative to the initial cluster count

The table provides a structured summary of dataset refinement across the pipeline and allows users to quantitatively evaluate how each screening criterion affects cluster quality. The table is generated based on the default pipeline order and is automatically updated if the sequence of pipeline steps is modified, allowing users to efficiently optimize the workflow for their specific recordings.

## DISCUSSION

SpikeCleaner provides a modular and interpretable framework for automated exclusion of clearly non-neuronal clusters from spike sorting outputs, addressing a critical intermediate step between automated clustering and expert manual curation. By combining rule-based classification with explicit reasoning labels and user-adjustable thresholds calibrated against manual annotations, the framework supports transparent and reproducible quality control across diverse electrophysiological datasets. Rather than replacing expert review, SpikeCleaner reduces the number of clusters requiring inspection and accelerates early-stage evaluation of spike sorting outputs, particularly during parameter optimization and dataset benchmarking. Should users determine that they are satisfied with SpikeCleaner, they can adopt its thresholds and decisions as conclusive thereby standardizing their pipelines to a greater extent than currently practiced in most scenarios.

It is important to note that other pipelines with similar goals already exist, namely BombCell^**14**^ and UnitRefine^**15**^. It is important to note that other high-quality tools with similar goals already exist, including BombCell^14^ and UnitRefine^15^. BombCell is available as a standalone MATLAB toolbox, with a more recently developed Python implementation and SpikeInterface integration. It can operate directly on Kilosort outputs and compute a range of waveform, spike-statistics, and autocorrelogram-based quality metrics, including estimates of refractory-period violations and unit contamination. Its metrics and unit labels can also be loaded into Phy for subsequent inspection and manual curation. UnitRefine is a machine-learning-based framework integrated with the SpikeInterface ecosystem that learns classification criteria from expert-labeled datasets. SpikeCleaner similarly operates downstream Kilosort and produces Phy-compatible outputs but uses an explicitly rule-based framework with user-adjustable thresholds, documented reasons for individual classification decisions, and tools for calibrating thresholds against laboratory-specific expert annotations.

A central feature of SpikeCleaner is the ability to adapt its classification thresholds to specific recording conditions and curation standards. Should users not choose to use our default parameters. This is important given the possible variability in datasets or recordings across varying species, labs, amplifiers or brain regions and does not force users to use a single set of criteria, although one such set is provided. This approach additionally does not require users to have *a priori* knowledge of thresholds but includes a tool that enables them to compare the outcomes of a series of systematically objective thresholds to their own expert judgments automatically - thereby enabling them to choose thresholds to match their own methods. It can enable custom thresholds that can be used for example within the confines of a given laboratory or dataset, and in a manner that is logged via parameter output files and is quantitative.

The use of hard-coded thresholds (even if re-defined by users) presents both weaknesses and strengths relative to training of AI-based systems to do classifications. AI-based classification may use features we as the developers never thought to quantify and therefore could produce better results in more scenarios. On the other hand, the thresholds used in any given run of SpikeCleaner are recorded and can therefore enable scientifically rigorous replication of procedures by other researchers and to date has produced high accuracy and other performance measures. Future work could directly compare the performance of SpikeCleaner to neural network-based tools formed around similar inputs and outputs.

A clear gap in our system, and many others, is the lack of development of cluster merge and split decisions. Such development would further reduce the manual burden associated with large-scale recordings and could be performed while preserving user oversight of final classification outcomes. However, split/merge represents a relatively orthogonal set of coding/decision-making tasks and is likely a harder problem to solve overall. We look forward to the development of such tools (or to their obviation by improving initial spikesorting algorithms), but that goal is not within the scope of this work.

Overall SpikeCleaner is a highly practical tool to either include or exclude neurons using either default parameters or to enable users to determine their own parameters. We are enthusiastic about the standardized and quantitatively based outputs this system can generate for a field with increasingly large datasets being used by broadening user base.

## CODE AVAILABILITY

Repository and Wiki: https://github.com/BrendonWatsonLab/SpikeCleaner

## ACKNOWLEDGMENTS

This work has been supported by funding from the NIH (MH107662, MH131592, NS141983, NS139668, MH131527), the Pritzker Neuropsychiatric Disorders Research Consortium and the University of Michigan Neuroscience Scholars fund to BOW.

## GENERATIVE AI CONTRIBUTIONS

Generative AI tools were used only for minor language editing and refinement of manuscript text. No AI tools were used for data analysis, interpretation, figure generation, or scientific decision-making.

## APPENDIX

### 1. Extraction of Waveforms

The output of **dz_filterWaveform()**, which is called within **dz_Curate():** animalname_filtered.mat file. Contains filtered ±2 ms waveform snippets and serves as the processed signal representation used for downstream feature extraction and quality control. Specifically, it stores several cluster-organized cell arrays: clippedWaveforms (number of clusters×1 cell), which contains the filtered waveform snippets for each cluster; meanWaveforms (number of clusters × 1 cell), which stores the average filtered waveform per cluster; bestWaveformsChannel (1×number of clusters cell), which records the selected best channel for each cluster based on waveform characteristics (e.g., maximum amplitude); bestWaveforms (number of clusters×1 cell), which contains the filtered representative waveform from the best channel; and channelCorrelations (number of clusters×1 cell), which stores inter-channel waveform correlation metrics used for spatial localization quality checks. Each cell corresponds to a specific cluster ID, allowing cluster-wise access to processed waveforms and derived metrics. By separating filtered waveforms into this dedicated .mat file, the pipeline avoids repeated filtering during threshold sweeps while ensuring that all subsequent feature computations (half-width, slope, amplitude, correlation-based QC) operate on consistently preprocessed spike-band signals.

**dz_Curate** internally calls **dz_extractWfVariables**, which first extracts a Three-Point Waveform of the maximum amplitude channel. To extract reliable waveform features, SpikeCleaner reduces each best-channel waveform (maximum amplitude waveform averaged over all the spikes in the cluster) to a physiologically meaningful three-point representation centered on the spike’s principal extremum. The best channel is defined as the site with the highest peak-to-trough amplitude.

*%finding the best polynomial fit for the best wf-just highpassed: to ignore the micro-fluctuations and obtain the global shape*

*[bestFittedValues]=dz_fitPolynomial(smoothenedbestwf,timeVector);*

*% Find peaks and troughs on smoothened and simply high-passed waveforms*

*[allLocsSorted, sortIdx] = dz_detectPeaksAndTroughs(bestFittedValues,timeVector); %%smoothened wf*

*[allLocsSorted1, sortIdx1] = dz_detectPeaksAndTroughs(bestwf,timeVector);%% from actual wf-max amplitude wf*

After detecting candidate peaks and troughs on both the smoothed and high-passed versions of the waveform, the algorithm removes redundant or near-duplicate extrema (e.g., adjacent peaks or troughs differing by <50 µV) and retains only the strongest.

The next step in selecting the 3-point waveform is to decide if it’s a positive spiking or a regular spiking neuron, which means checking if the peak is more pronounced or the trough is deeper, and making sure that the deepest trough has peaks on either side, Regular Spiking, or the highest peak has troughs on either side, Positive Spiking.

~~~
*
[maxPeak, peakIdx] = max(sortIdx1); % Find the largest peak
[minTrough, troughIdx] = min(sortIdx1); % Find the deepest trough
if abs(minTrough) > abs(maxPeak)
 spikingType = ‘Regular Spiking’;% More negative trough : Regular Spiking
 if (troughIdx > 1 && troughIdx < length(sortIdx1)) %Ensure the deepest trough has peaks on both sides
   if sortIdx1(troughIdx - 1) > 0 && sortIdx1(troughIdx) < 0 && sortIdx1(troughIdx + 1) > 0
    selectedIndices = [troughIdx - 1, troughIdx, troughIdx + 1]; % Peak-Trough-Peak
    end
else
   sym = 1; % Noise : if even after being regular spiking it has less than 3 indices
end
else
 spikingType = ‘Positive Spiking’;
 % Only do this in the Positive Spiking condition before setting selectedIndices
 if (∼isempty(baselinePeak) && peakIdx==1)
   %in case, because of consecutive peak removal, the highest peak is at 1, in positive spiking
   % Prepend baseline peak and loc so it appears before the main peak: as in positive spiking, there is always a
   % baseline peak and not a flat baseline before the main peak. So we need to conserve it and prepend it. sortIdx1(peakIdx+2)=[];
   allLocsSorted1(peakIdx+2)=[];
   sortIdx1 = [baselinePeak, sortIdx1];
   allLocsSorted1 = [baselineLoc, allLocsSorted1];
   % Since we inserted a new point at the beginning, we shift peakIdx by +1
   peakIdx = peakIdx + 1;
end
 selectedIndices = [peakIdx - 1, peakIdx, peakIdx + 1]; %trough-peak-trough
end
*
~~~

Based on the 3-point waveform extracted, it further calculates the local set of channels that a user wants to look at, for example, the 8 closest channels to the maximum amplitude channel. For this set of channels, we extract the difference in amplitude from the maximum amplitude channel. We also extract halfwidth, slope, and the spike type (regular/positive) of the maximum amplitude channel for every cluster.

To calculate the half-amplitude level between the waveform’s peak and trough we locate the time points on the left and right sides of the main extremum where the waveform crosses this half-amplitude value. The time difference between these two crossing points defines the spike half-width. Using these same points, we also calculate the rising and falling slopes of the waveform by measuring how quickly the signal changes between the extremum and each half-amplitude crossing. These slope measurements help assess whether the waveform has physiologically realistic spike dynamics.

We do this beforehand, so that no matter what pipeline flow user chooses, we have all the variables for all the tests available.

~~~
*
halfAmplitude = ypoint;
halfAmplitudeIndex=xpoint;
[∼, leftIdx] = min(abs(bestwf(allLocsSorted_idx(1):allLocsSorted_idx(2)) - halfAmplitude));
leftIdx = leftIdx + allLocsSorted_idx(1) - 1; % Adjust index to global waveform
[∼, rightIdx] = min(abs(bestwf(allLocsSorted_idx(2):allLocsSorted_idx(3)) - halfAmplitude));
rightIdx = rightIdx + allLocsSorted_idx(2) - 1; % Adjust index to global waveform
halfWidth = timeVector(rightIdx) - timeVector(leftIdx);
Slope1 =(bestwf(allLocsSorted_idx(2)) - bestwf(leftIdx)) / …
   (timeVector(allLocsSorted_idx(2)) - timeVector(leftIdx) + eps);
Slope2 =(bestwf(rightIdx) - bestwf(allLocsSorted_idx(2))) / …
   (timeVector(rightIdx) - timeVector(allLocsSorted_idx(2)) + eps);
*
~~~

### 2. Extraction of ACG & ACG Proportions

**dz_Curate()** calls an internally calls dz_extractAcgVariables which in turn calls **dz_autoCorr()**. ACG is computed by measuring how often spikes from a single neuron occur at specific time intervals relative to other spikes from the same neuron. A symmetric time window is defined around zero lag (for example, −25 ms to +25 ms) and divided into equal-sized bins (e.g., 1 ms bins). For each spike in the spike train, the time differences between that spike and all other spikes are calculated. The self-comparison at zero lag is excluded to avoid artificially inflating the center bin. These time differences represent how far before or after other spikes occurred relative to the reference spike. The differences are then sorted into the predefined time bins using a histogram. This process is repeated for every spike, and the histograms are summed across all reference spikes. The result is a distribution showing how frequently spikes occur at different time lags relative to one another, centered at zero. It returns all the bin values of the number of spike pairs separated by that time difference in **currentacg** and bin centers **in binCenters**, which represent time offsets around 0. We then shift the centers to make sure the center bin is at 0, because MATLAB calculates bin centers using floating-point arithmetic. Because of rounding precision, the center bin might not be exactly 0. This would guarantee that the ACG is precisely centered at 0 lag and avoids floating-point misalignment errors. Here, we also extract the proportion fill of the center of the ACG with respect to the shoulders, and whether most of the ACG bins are empty or not.

ACG Extraction: t1&t2 are the spike trains of the same cluster.

~~~
*
numBins=25;
binSize= 0.001;%1ms
total=(2*numBins)+1;
half=(total-1)/2;
 edges = (-half:half + 1) * binSize;
 % Iterate over spikes in t1 to compute ACG
   for i = 1:length(t1)
    diffs = t2(t2 ∼= t1(i)) - t1(i); % Excludes self-comparison
    cch = cch + histcounts(diffs, edges);
 end
*
~~~

It computes **ACG center-to-shoulder proportions**, i.e., how large the spike counts near **zero lag** are compared to the **shoulder peaks** of the autocorrelogram. This helps detect **refractory period violations**, which indicate multi-unit activity (MUA) rather than single units. The shoulder peak of the ACG is estimated by first excluding bins around zero lag to avoid the refractory period region, and then searching for prominent peaks in the surrounding shoulder region. Local maxima are detected using a peak-finding algorithm with minimum prominence and spacing constraints to reduce noise effects. The largest three peaks are selected and averaged to obtain a stable estimate of the shoulder peak amplitude. This average shoulder value represents typical spike timing structure outside the refractory window and is used as a reference to normalize central ACG bins when assessing refractory period violations and identifying multi-unit activity.

~~~
*
% Find first peak after central refractory zone
searchStart = zeroLagIndex + 3; % skip 3 bins on right side of center
seg = currentacg(searchStart:end);
%%finding the highest peak
 k =20;
 if numel(seg) > k && all(seg(1) > seg(2:k+1))
  peakRel = 1;
  peakVal = seg(1)
 else
  [pks, locs] = findpeaks(seg, …
  ‘MinPeakProminence’, max(5, 0.02*max(seg)), …
   ‘MinPeakDistance’, 2);
  if isempty(pks)
   [peakVal, peakRel] = max(seg); % fallback
  else
   [sortedPks, idx] = sort(pks, ‘descend’);
   % Number of peaks to average
   n = min(3, length(sortedPks));
   % Take top n peaks
   topPeaks = sortedPks(1:n);
   % Average them
   peakVal = mean(topPeaks);
  end
end
proportion=cchzero/peakVal; %proportion of center peaks in comparison to the shoulder of the acg
*
~~~

IsEmptyCheck:

This step checks whether the ACG for a cluster is largely empty, which can indicate insufficient spike counts or unreliable temporal structure. It computes the proportion of bins in the ACG that contain zero values, and if more than 50% of the bins are empty, the cluster is flagged as having an empty ACG. In such cases, no center-to-shoulder proportion is calculated, and the proportion values are set to NaN before moving to the next cluster. This prevents unreliable ACG-based metrics from influencing later classification steps.

~~~
*
%% check if empty 15 bins on either side
zeroLagIndex = ceil(length(currentacg) / 2);
fracZero = sum(currentacg == 0) / length(currentacg);
if fracZero > 0.5
  isEmpty(ix) = true;
  proportion= nan(1,6);
  Proportions{ix}=proportion; % saving it in the cell array
  continue;
end
*
~~~

### 3. Building a Pipeline Flow for Quality Check Points that a User can Rearrange

SpikeCleaner works based on consecutive quality checks and the flow and entry points to those checks are specified by the flow of the Pipeline. Based on the flow of the pipeline that the user chose, SpikeCleaner sends all units consecutively to the respective quality tests. All units start as ‘Good/Single Units’ and after each test are re-labelled with the updated results. The units that have been discarded/labelled as Noise/MUA in the previous step do not proceed to the next step and are skipped over. This ensures the reason for eviction matches the flow of the pipeline and avoids redundant processing of units.

Default Pipeline: l*owFiring’,’amplitude’,’halfWidth’,’slope’,’correlation’,’acgEmpty’,’acgAll’,’mua’*

*pipeline={‘lowFiring’,’amplitude’,’halfWidth’,’slope’,’correlation’,’acgEmpty’,’acgAll’,’mua’};*

*dz_classifyAllUnits(‘mode’,’strict’,’maxHW’,0.8,’minAmp’,50…,’pipeline’=pipeline)*

The SpikeCleaner pipeline is modular and user configurable. The order of quality-control steps is defined by a user-provided pipeline list, and the algorithm executes each step sequentially using a switch–case structure. At each iteration, the pipeline reads the next step name (e.g., *lowFiring, amplitude, halfWidth, slope, correlation, ACG checks*, or *MUA detection*) and applies the corresponding evaluation function. Users can modify the sequence, remove steps, or insert additional checks by changing the pipeline list, allowing flexible control over the order in which clusters are filtered or classified.

~~~
*
for k = 1:length(pipeline)
  step = pipeline{k};
  switch step
   case ‘lowFiring’
     dz_evaluateLowFiringRates(recordingDuration, firingThreshold);
     radarLabels{k} = ‘Low Firing’;
   case ‘correlation’
     dz_analyzeCorrelation(wfs, channelCorrelations,wfdata,pos,nLocalChannels);
      radarLabels{k} = ‘Correlation’;
   …
   …
end
*
~~~

